# A longitudinal model of human neuronal differentiation for functional investigation of schizophrenia polygenic risk

**DOI:** 10.1101/211581

**Authors:** Anil P.S. Ori, Merel H.M. Bot, Remco T. Molenhuis, Loes M. Olde Loohuis, Roel A. Ophoff

## Abstract

There is a pressing need for *in vitro* experimental systems that allow for interrogation of polygenic psychiatric disease risk to study the underlying biological mechanisms. We developed an analytical framework that integrates genome-wide disease risk from GWAS with longitudinal *in vitro* gene expression profiles of human neuronal differentiation. We demonstrate that the cumulative impact of risk loci of specific psychiatric disorders is significantly associated with genes that are differentially expressed across neuronal differentiation. We find significant evidence for schizophrenia, which is driven by a longitudinal synaptic gene cluster that is upregulated during differentiation. Our findings reveal that *in vitro* neuronal differentiation can be used to translate the polygenic architecture of schizophrenia to biologically relevant pathways that can be modeled in an experimental system. Overall, this work emphasizes the use of longitudinal *in vitro* transcriptomic signatures as a cellular readout and the application to the genetics of complex traits.

## Introduction

Major psychiatric disorders feature a high heritability but have a largely unknown etiology^1,2^. The increasing sample sizes of genome-wide association studies (GWAS) successfully result in identification of more susceptibility loci for these disorders^3^. A major challenge is to understand and interpret the cumulative impact of many loci that collectively contribute to psychiatric disease risk and how to translate this complex polygenic architecture to biological pathways that drive the underlying molecular and cellular disease processes. Lack of applicable *in vitro* model systems and a framework to study polygenic psychiatric risk hinders the translation of genetics findings to disease biology^4^.

Early brain development has been implicated in psychiatric disorders such as schizophrenia (SCZ)^5–8^, autism spectrum disorder (ASD)^9, 10^, and self-reported depression (SRD)^11^. Differentiation of human embryonic stem cells (hESCs) into neuronal lineages has been demonstrated to hold great promise to model early brain development^12–14^, and may thus offer a unique opportunity to study psychiatric disease biology *in vitro.* However, it has remained unclear whether the molecular dynamics underlying *in vitro* human neuronal differentiation are associated with polygenic psychiatric disease susceptibility.

We set out to investigate *in vitro* human neuronal differentiation in the context of polygenic psychiatric disease risk. To accomplish this, we performed a densely-sampled time series experiment and robustly detected transcriptome-wide changes across neuronal differentiation. To study the aggregate impact of risk loci, we integrated longitudinal *in vitro* gene expression signatures with GWAS summary statistics of major psychiatric disorders. We observe significant enrichment of genetic risk for multiple disorders in genes that are upregulated across differentiation. We further show that this effect is strongest for SCZ and primarily driven by a longitudinal gene cluster that is involved in synaptic functioning. These findings support to use of *in vitro* neuronal differentiation as a promising model system to study genetic psychiatric risk, particularly in the context of schizophrenia.

## Material and Methods

### Approval for stem cell research

This study and all described work was approved by the University of California, Los Angeles Embryonic Stem Cell Research Oversight (ESCRO) committee.

### In vitro human neuronal differentiation

WA09(H9)-derived hNSCs were commercially obtained (Gibco) as neural progenitors and subsequently expanded as adherent culture according to the manufacturer’s guidelines (Supplementary Methods). Low passage hNSCs (< 4 passage rounds) were plated in 12-well plates coated with poly-D-lysine (0.1 mg/mL, VWR) and laminin (4.52ug/cm^2^, Corning^™^) at 1.5×10^5^ cells, which were equally distributed and subsequently cultured in expansion medium as described above. After 24h of proliferation, media was changed to neuronal differentiation medium consisting of Neurobasal^®^ Medium (Gibco), 2% B-27^®^ Serum-Free Supplement (Gibco), 2mM GlutaMax™-I Supplement, 0.05 mM β-mercaptoethanol (Gibco), and 1x Pen Strep. Media was changed every 2-3 days.

### Experimental design and assessment of gene expression

Human neural stem cells were differentiated over a course of 30 days and RNA harvested at seven time points (day 0, 2, 5, 10, 15, 20, and 30) in triplicates or quadruplicates (n = 24) (Supplementary Methods). Genome-wide array-based transcriptome data was collected at the UCLA Neuroscience Genomics Core using Illumina’s HumanHT-12 v4 Expression BeadChip Kit.

### Data preprocessing and quality control

Gene expression data was extracted using the Gene Expression Module in GenomeStudio Software 2011.1. Data was background corrected with subsequent variance-stabilizing transformation and robust spline normalization was applied^15, 16^. We excluded low quality probes and subsequently performed sample outlier detection by Euclidean distance and standardized connectivity (see Supplementary Methods). The FactoMineR package (v1.28) in R was used to perform principal component analysis (PCA). For subsequent downstream analyses, we used the normalized expression values of 19,012 high quality filtered probes for all 24 samples.

### Transcriptome-based in vitro cellular identity

We identified cell-type specific genes of neurons, astrocytes, oligodendrocyte precursor cells (OPC), newly formed oligodendrocytes (NFO), myelinating oligodendrocytes (MO), microglia, and endothelial cells from mouse cerebral cortex^17^ (Supplementary Methods). Next, we extracted normalized gene expression values of these genes for each cell type from our own *in vitro* dataset. We then standardized expression values to time point zero and calculated mean standardized expression levels of cell type-specific genes for these seven cell types across time points. This allowed us to investigate *in vitro* cellular identity across differentiation using cell-type specific gene expression information of many genes for these seven cell types.

### Transition mapping to a spatiotemporal atlas of early human brain development

To investigate global transcriptomic matching between *in vitro* gene expression profiles and *in vivo* gene expression profiles of neocortical brain regions, we applied transition mapping (TMAP), which is implemented in the online CoNTExT bioinformatic pipeline (https://context.semel.ucla.edu)^14^. Analyses were run for *in vitro* time points day-0 vs day-30, day-0 vs day-5, day-5 vs day-15, and day-15 vs day-30 across both temporal and spatial dimensions of human cortical development (see Supplementary Methods).

### Time-series differential gene expression analysis

Two multivariate empirical Bayes models were used to identify differentially expressed genes across *in vitro* neuronal differentiation. The first method was implemented in the Timecourse package (v 1.42) in R. We used the mb.long() function to calculate the one-sample T^*2^ statistic that ranks genes based on their log10 probability to have differential expression over time^18^. Bayesian Estimation of Temporal Regulation (BETR), an extension of the first approach, uses a flexible random-effect model that allows for correlations between the magnitude of differential expression at different time points^19^. BETR is implemented in the betr package (v 1.26) in R. Differentially expressed genes were classified as the union of the set of genes with a probability of 1.0 using BETR and an equally-sized set of top ranked genes using the T^2^-statistic.

### Fuzzy c-means cluster analysis

To identify probes with similar expression patterns across differentiation, we applied fuzzy c-means clustering to all differentially expressed probes. We calculated cluster membership values using the fclusList() and membership() function in the Mfuzz package in R^20, 21^. See Supplementary Methods for more details. Each probe receives a membership value for each cluster. Probe membership values represent gene affiliation to a cluster and highlights the extent of similarity in expression between genes. These values were used for subsequent downstream analyses. We annotated clusters using Database for Annotation, Visualization, and Integrated Discovery (DAVID, v6.8) and probes with a membership > 0.5 (Supplementary Methods).

### Integration of GWAS data with in vitro transcriptomic signatures

We first mapped Illumina probe IDs to Ensembl gene IDs using NCBI build 37.3, removed duplicate Ensemble IDs, and extended gene boundaries symmetrically by 10kb to include regulatory regions. Probe T^2^-statistic and cluster membership values were collapsed per gene ID using the mean value across probes. The mean gene-level T^2^-statistic was then log-transformed and the mean cluster membership values rank-transformed. These mean gene values were then used to integrate *in vitro* signatures with GWAS data and to study the cumulative impact across risk loci using Multi-marker Analysis of GenoMic Annotation (MAGMA) and stratified LD score regression (sLDSR).

### GWAS summary statistics and ancestry matched reference panels

GWAS summary statistics were obtained for SCZ^22^, major depressive disorder (MDD)^23^, SRD^11^, bipolar disorder (BPD)^24^, ASD^25^, attention deficit hyperactivity disorder (ADHD)^26^, cross disorder^27^, Alzheimer’s disease (AD)^28^, and adult human height^29^ (Supplementary Methods and Table S2). For each trait we used the most recent GWAS summary statistics that was publically available at the time of the analysis. The 1000 Genomes Project Phase 3 release (1KG) was used as reference panel to model LD^30^. We used 503 individuals of European ancestry and 301 individuals of East Asian ancestry in analyses of GWAS data derived from target population of Europeans and Han Chinese, respectively.

### MAGMA gene-set analysis

Multi-marker Analysis of GenoMic Annotation (MAGMA v1.06)^31^ was used to run “gene property” analyses, which uses a multiple regression framework to associate a continuous gene variable to GWAS gene level p-values. SNPs were mapped to genes using Ensembl gene IDs and NCBI build 37.3 gene boundaries +/− 10kb extensions using the --annotate flag. For each phenotype, we generated gene-level p-values by computing the mean SNP association using the default gene model (‘snp-wise=mean’). We only included SNP with minor allele frequency (MAF) > 5% and dropped synonymous or duplicate SNPs after the first entry (‘synonym-dup=drop-dup’). For each annotation, we then regressed gene-level GWAS test statistics on the corresponding gene annotation variable using the ‘--gene-covar’ function while adjusting for gene size, SNP density, and LD-induced correlations (‘--model correct=all’), which is estimated from an ancestry-matched 1KG reference panel. In all analyses, we included only genes for which we had both the gene variable and GWAS gene level test statistic available. Testing only for a positive association, i.e. enrichment of GWAS signal, we report one-sided p-values along with the corresponding regression coefficient.

### Stratified LD Score Regression

We applied a recent extension to stratified LD score regression (sLDSR), a statistical method that partitions SNP-based heritability (h^2^) from GWAS summary statistics^8^. This extension allows us to quantify the effects of continuous-valued annotations on the heritability^32^. For each annotation, we first estimated partitioned LD scores using the ldsc.py --l2 function with MAF > 5%, a 1 centimorgan (cm) window, and an ancestry-match 1KG reference panel (Supplementary Methods). We ran sLDSR (ldsc.py --h2) for each annotation of interest while accounting for the full baseline model, as recommended by the developers^8, 32^, and an extra annotation of all genes detected in our *in vitro* model (n = 12,414). That is, for each annotation we ran the following model;

1. Full baseline model with 53 annotations.
2. Annotation of all genes detected during *in vitro* neuronal differentiation.
3. Annotation of interest (e.g. cluster membership).

If an annotation of interest (3) is associated with increased h^2^, LD to SNPs with large values of that annotation will increase the *χ*^2^ statistic of a SNP more than LD to SNPs with smaller values. To determine if this effect is significant and specific to this annotation, it estimates the contribution of that annotation to the per-SNP h^2^ while accounting for the baseline and the all genes detected annotation (1 + 2). As we only test for a positive association, we report the contribution to the per-SNP h^2^ (τ) and the associated one-sided p-value, which is calculated using standard errors that are obtained via a block jackknife procedure^8, 33^.

### Statistical analyses

Statistical analyses were performed with R (https://www.r-project.org) or an otherwise specified algorithm. Significance with MAGMA and sLDSR was determined at a one-sided α level of 0.05. If applicable, Bonferroni correction for multiple comparison is applied and denoted in figures and tables. For all analyses, further details on statistics are available in the supplementary results.

## Results

### Longitudinal in vitro gene expression profiling confirms neuron-specific differentiation and matches in vivo human cortical development

To study the molecular dynamics underlying *in vitro* human neuronal differentiation, we differentiated an hNSC line (WA09/H9) to a neuronal lineage across 30 days. Genome-wide gene expression profiles were assayed densely at seven time points in at least triplicates (n=24 samples). Principal component analysis (PCA) on normalized gene expression values shows a large proportion of the variance in expression to be explained by the differentiation process, with minimal effects of technical variation (Figure 1A & S1). Investigation of transcriptome-based cell type-specific gene expression signatures of major classes of cell types in the cerebral cortex shows that relative neuronal gene expression increases as neuronal differentiation progresses over time (Figure 1B). There is no evidence of glial- or endothelial-specific gene expression, which confirms a broadly neuronal *in vitro* cellular identity.

**Figure 1.**
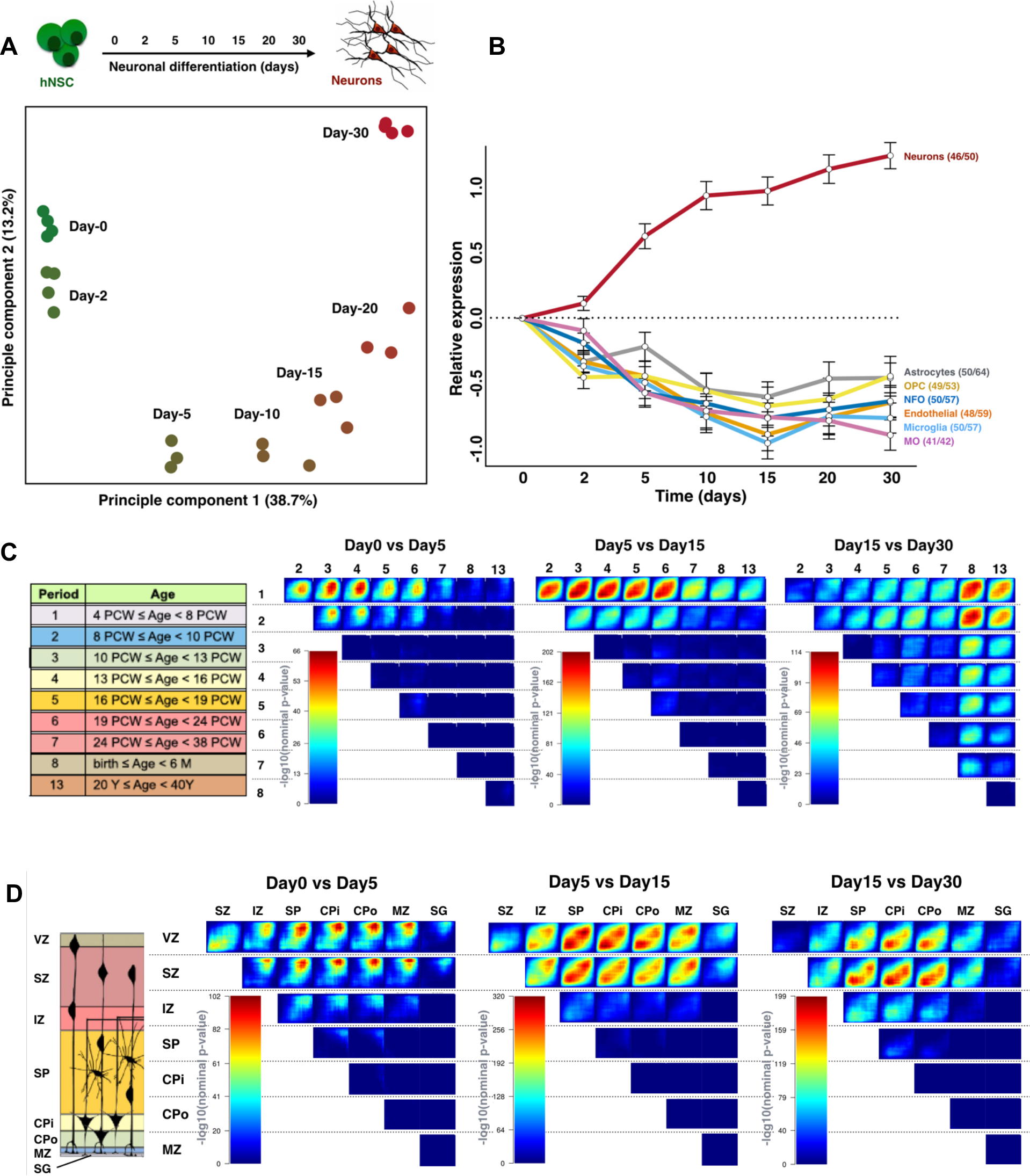
*In vitro* gene expression profiles confirm a neuron-specific differentiation process and match *in vivo* human cortical development. (A) PCA of *in vitro* transcriptomic data with PC1 (x-axis) and PC2 (y-axis) visualized. Variance explained per component is shown in parentheses. Time points are color-coded and labeled by days across differentiation. (B) Transcriptome-based cellular identity is shown by average expression of cell type specific genes across days of differentiation. Cell types are highlighted by their name and corresponding color. The first number in the parentheses represents the number of genes for which the average expression is plotted. The second number represents the corresponding number of probes assayed. OPC = oligodendrocyte precursor cells, NFO = newly formed oligodendrocytes, MP = myelinating oligodendrocytes. (C+D) TMAP output visualizes the amount of overlap between *in vitro* and *in vivo* DGE profiles colored by −log10(p-value) (see figure S2 for more details on interpretation). Abbreviations and numbering above maps correspond to schematic representations on the left (adopted from Stein et al., 2014) of different developmental stages (C) and laminae (D). VZ = ventricular zone, SZ = subventricular zone, IZ = intermediate zone, SP=subplate zone, CPi= inner cortical plate, CPo = outer cortical plate, MZ = marginal zone, PCW = post conception weeks, M = months, Y = years, Period = developmental stage.

Having established that the *in vitro* differentiation process is predominantly neuronal, we applied transition mapping (TMAP) to assess the correspondence of longitudinal *in vitro* transcriptome data to *in vivo* signatures of human cortical development. TMAP uses a spatiotemporal transcriptome atlas of the human neocortex and laminar expression data to assess global overlap in differential gene expression (DGE) profiles between *in vitro* time points and *in vivo* brain developmental stages or laminae of the human neocortex. We find significant matching between the *in vitro* longitudinal DGE profiles (day-0 vs day-30) and *in vivo* developmental stage from 4 weeks post-conception (PCW) to 24 PCW (Figure S2). This overlaps with the primary period of neurogenesis in the neocortex, which starts around 6 PCW^34, 35^. To gain more insight into this overlap, we partitioned the TMAP analyses in three comparisons and examined how *in vitro* to *in vivo* matching progressed over time across *in vitro* neuronal differentiation. We see a clear progression in matching from early developmental stages to later stages (Figure 1C). For example, *in vitro* day-0 vs day-5 show strong overlap with *in vivo* period-1 (4-8 PCW) vs period-4 (13-16 PCW), while *in vitro* day-15 vs day-30 shows stronger overlap with *in vivo* period-2 (8-10 PCW) vs period-8 (birth-6M). Similarly, *in vitro* longitudinal DGE shows progression from overlap of early time points with inner laminae, to overlap with more upper cortical layers as *in vitro* neuronal differentiation advances (Figure 1D and S2).

### In vitro neuronal differentiation reveals specific longitudinal gene clusters

To identify biological pathways associated with neuronal differentiation, we applied an analysis framework specifically tailored to time-series gene expression data (see Methods and Supplementary Methods). A total of 7,734 probes, mapping to 5,818 genes, were differentially expressed over time (Figure S3). We find that these genes are, on average, more constrained to genetic variation compared to non-differentially expressed genes (Supplementary Results). Using only differentially expressed probes, we next applied fuzzy c-means clustering and identified eight distinct longitudinal gene clusters (Figure 2 and S4). For each probe, we generated a corresponding cluster membership value, representing the degree to which a gene belongs to a cluster. To identify most informative biological interpretation of each cluster, we analyzed genes with high cluster membership for enrichment of functional annotations using DAVID (Supplementary Methods and Table S1).

**Figure 2.**
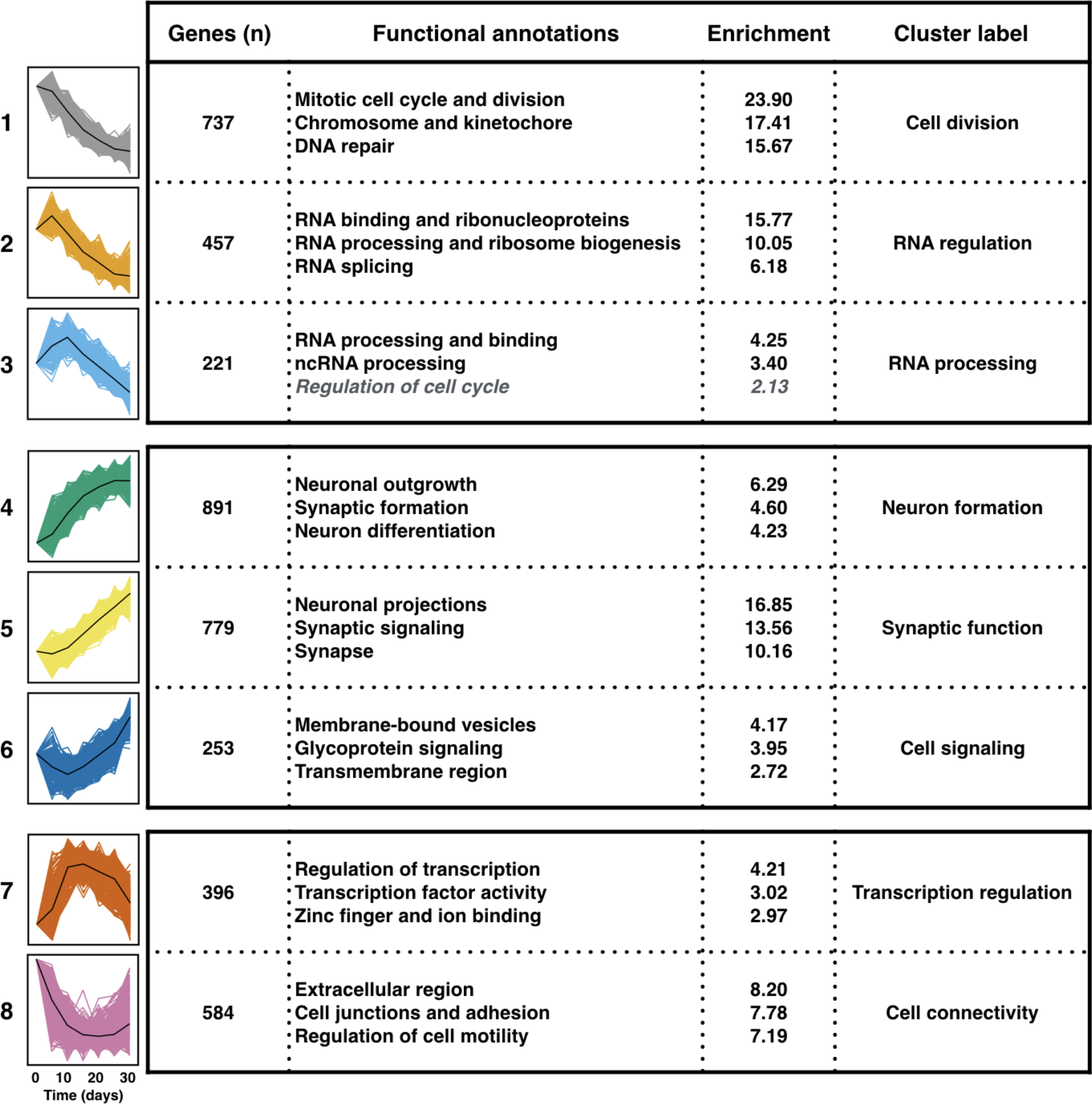
Identified gene clusters highlight biological pathways important for neuronal differentiation. Significant functional annotations and corresponding enrichment score are shown for each gene cluster. Longitudinal gene expression is visualized for high member genes only (black line represents mean gene expression). Each cluster is color-coded with the number of genes at membership > 0.5 denoted. See table S1 for full annotation results.

We identified three clusters with decreasing gene expression over time that are significantly enriched for cell division and RNA regulation and processing genes, reflective of stem cell proliferation and cell fate determination that is tightly controlled and regulated by RNA dependent processes^36^. Second, there are three clusters showing increased gene expression levels over time that are primarily enriched for neuronal processes, such as neuron formation and synaptic function. Another independent cluster shows an inverted U-shaped expression pattern during development, enriched for genes involved in transcriptional regulation. The final cluster is enriched for genes involved in extracellular region and cell adhesions. These processes are important for cell connectivity and have also been implicated in cell proliferation and neuronal migration^37, 38^. Together, these eight gene clusters reveal different biological mechanisms that are associated with neuronal differentiation and consistent with known biology of neurodevelopment. We hypothesize that the study of these longitudinal gene expression clusters can help decipher disease mechanisms involved in psychiatric phenotypes.

### Differentially expressed genes are enriched for polygenic psychiatric disease risk

To examine how aggregate psychiatric disease risk is distributed across genes that are important for neuronal differentiation, we applied gene-set analysis and partitioning of h^2^ with MAGMA and sLDSR, respectively. We investigated the cumulative impact of risk loci associated with major psychiatric disorders using GWAS results from large-scale studies. See Supplementary Table S2 for details on all included phenotypes. For MDD, we included GWAS results from the China Oxford and VCU Experimental Research on Genetic Epidemiology (CONVERGE) consortium^23^ and 23andMe Inc., a personal genetics company^11^. The latter uses SRD as a proxy for major depression. Alzheimer’s disease (AD) and adult human height served as non-psychiatric control phenotypes that are heritable and polygenic. We used a two-step approach where we first investigated disease susceptibility on overall differential expression level and subsequently proceeded to deconstruct these associations across the longitudinal gene clusters. Together, these analyses allow us to integrate *in vitro* longitudinal transcriptomic signatures with GWAS summary statistics and assess if our approach can capture the polygenic nature of psychiatric disorders.

We find that genes that are differentially expressed across *in vitro* neuronal differentiation are enriched for genetic risk of multiple psychiatric disorders. We find significant effects with MAGMA for SCZ (P=0.001), ADHD (P=0.002), and SRD (P=0.003) (Table 1 and Table S3). With sLDSR, we find nominally significant effects for SCZ (P=0.01) and SRD (P=0.02) and a suggestive association for ADHD (P=0.06) (Table 1 and Table S4). We observed a suggestive enrichment for BPD, and no enrichment for the cross disorder, ASD, MDD CONVERGE or for adult height and AD.

**Table 1.** Differentially expressed genes are enriched for polygenic risk of multiple psychiatric disorders. Shown are results of MAGMA and sLDSR for differentially expressed genes. P-values highlighted in bold show phenotypes that survive multiple testing correction (n=9). See Table S3 and S4 for more details. Beta = regression coefficient, SE = standard error, Beta_std = change in Z-value given a change of one standard deviation in log T2 statistic, *τ* (tau) = the contribution to the per-SNP h^2^.

| Phenotype | MAGMA |  |  | sLDSC |  |
| --- | --- | --- | --- | --- | --- |
| | Beta (SE) | Beta_std | P-value | $\tau$ (SE) | P-value |
| **Psychiatric** |  |  |  |  |  |
| Schizophrenia | 0.022 (0.007) | 0.094 | **0.001** | $1.70 \times 10^{-9}$ ($7.45 \times 10^{-10}$) | 0.01 |
| ADHD | 0.014 (0.005) | 0.059 | **0.002** | $1.92 \times 10^{-9}$ ($1.25 \times 10^{-9}$) | 0.06 |
| Self-reported depression | 0.013 (0.005) | 0.057 | **0.003** | $4.34 \times 10^{-10}$ ($2.10 \times 10^{-10}$) | 0.02 |
| Bipolar disorder | 0.007 (0.005) | 0.032 | 0.06 | $6.16 \times 10^{-9}$ ($3.64 \times 10^{-9}$) | 0.05 |
| Cross disorder | 0.005 (0.005) | 0.020 | 0.16 | $1.19 \times 10^{-9}$ ($1.00 \times 10^{-9}$) | 0.12 |
| MDD CONVERGE | 0.000 (0.004) | -0.001 | 0.51 | $6.07 \times 10^{-9}$ ($4.39 \times 10^{-9}$) | 0.08 |
| ASD | 0.000 (0.004) | -0.002 | 0.54 | $2.97 \times 10^{-9}$ ($3.48 \times 10^{-9}$) | 0.20 |
| **Neurodegenerative** |  |  |  |  |  |
| Alzheimer's disease | 0.003 (0.004) | 0.015 | 0.22 | $1.30 \times 10^{-10}$ ($1.02 \times 10^{-9}$) | 0.45 |
| **Non-brain** |  |  |  |  |  |
| Height | 0.009 (0.011) | 0.037 | 0.21 | $-1.62 \times 10^{-9}$ ($1.36 \times 10^{-9}$) | 0.88 |

We next investigated whether enrichment across differentially expressed genes was driven by up- or downregulation of genes during differentiation. For SCZ, we find that the effect is driven by genes that are upregulated (MAGMA P=5.0×10^−7^, sLDSR P=6.1×10^−5^) and not by genes that are downregulated (MAGMA P=0.98, sLDSR P=0.61) (Figure 3 and Figure S5). For SRD, we only find a stronger enrichment in upregulated genes with MAGMA (P=3.5×10^−4^), while ADHD shows no specific evidence for either up or downregulated genes.

**Figure 3.**
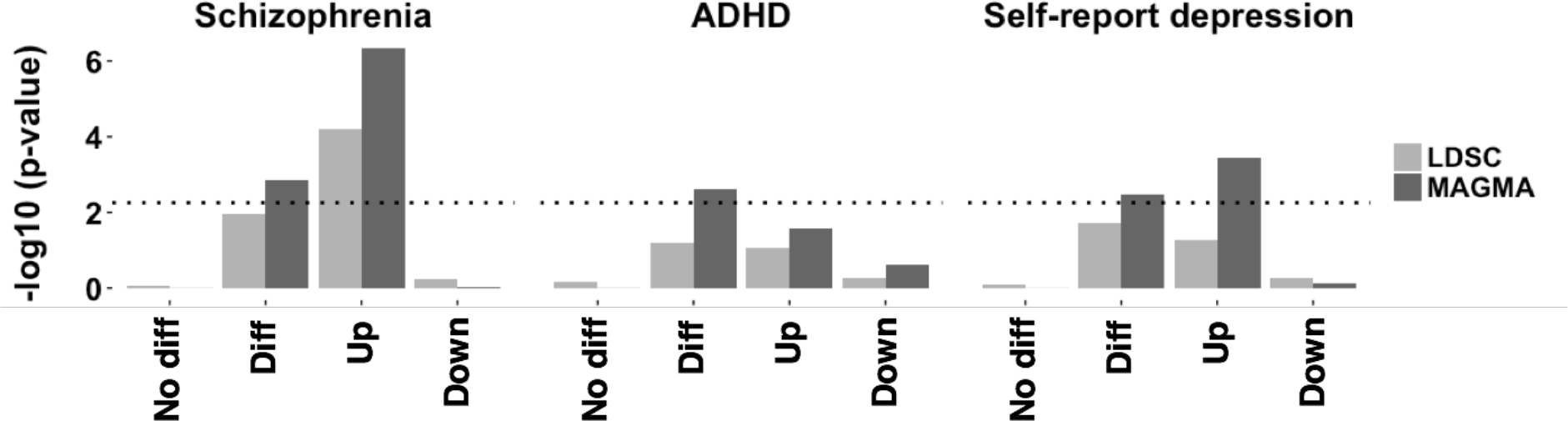
Schizophrenia polygenic risk lies in genes up-regulated during neuronal differentiation. A more detailed investigation of the effect of differentially expressed genes on the heritability of SCZ, ADHD, and SRD. The y-axis denotes the −log10 P-value of the enrichment. No diff = genes that are not differentially expressed; Diff = log (T^2^-statistic) as shown in Table 1; Up = genes up-regulated during differentiation; Down = genes down-regulated during differentiation. The dotted line represents the threshold for P=0.0056 (n=9 traits).

### Psychiatric disease risk aggregates to specific longitudinal gene clusters

Next, we explored the relationship between differentially expressed genes and disease risk on cluster level. For this analysis, we only included traits that show significant disease enrichment across differentially expressed genes using MAGMA after correcting for multiple testing (SCZ, ADHD, SRD) and our control traits (AD, height). These disease traits showed at least a nominally significant effect with sLDSR as well. Using both MAGMA and sLDSR, we integrated cluster membership values with GWAS summary statistics (n=5) and assessed whether genome-wide disease risk aggregates to any of the eight experimentally identified longitudinal gene clusters. Overall, MAGMA and sLDSR show a strong concordance across phenotypes and clusters (rho = 0.92, p<2.2×10^−16^, n=40, see also Figure S6). After Bonferroni correction (n=40), we find five significant phenotype-cluster associations with MAGMA and three with sLDSR (Figure 4 and Table S5/S6).

**Figure 4.**
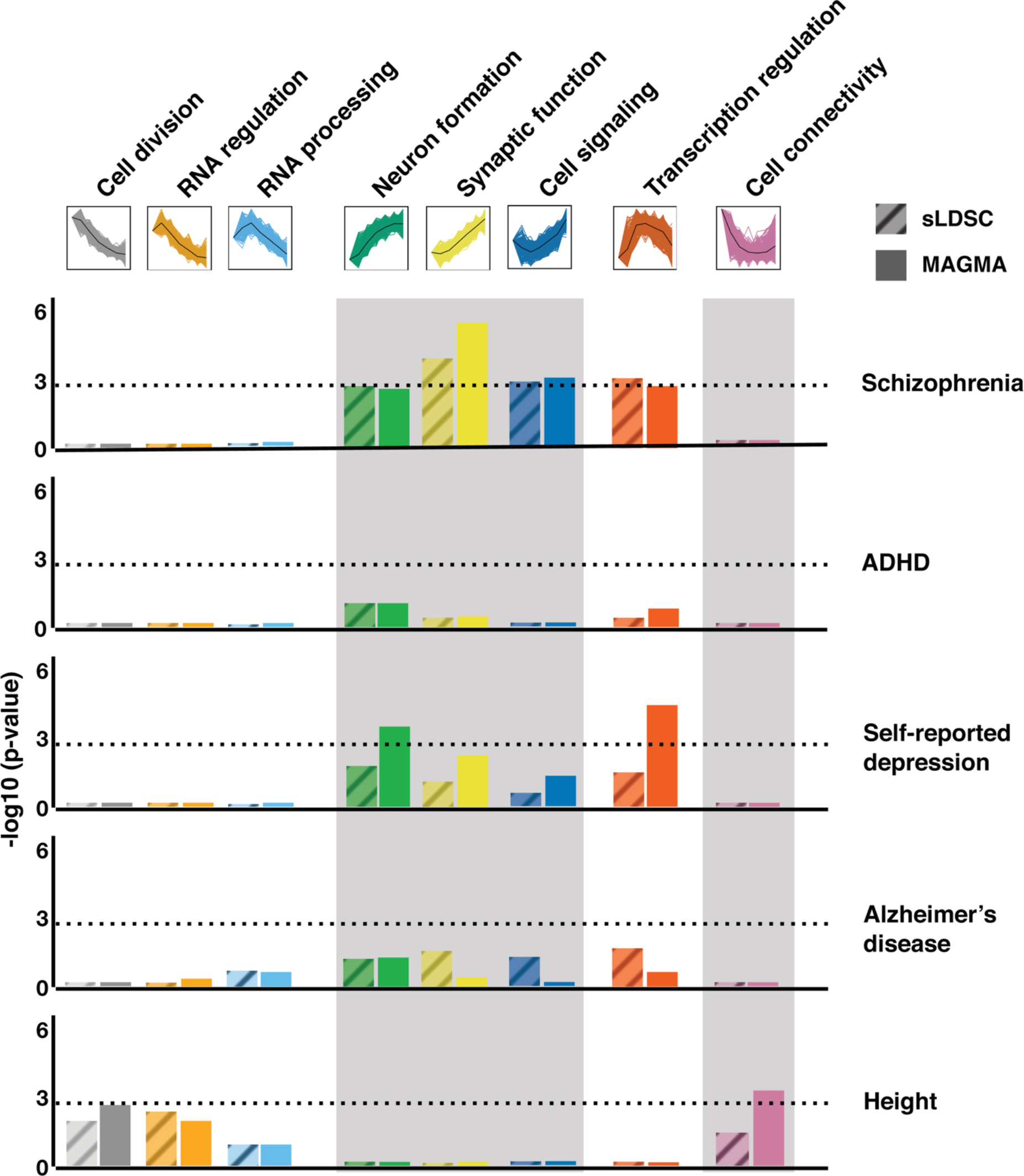
Polygenic psychiatric risk is distributed across specific longitudinal gene clusters. Results from sLDSC (diagonal pattern) and MAGMA (solid colors) are shown for each phenotype (labels on the right) colored by gene cluster. Gene cluster annotation and cluster expression pattern are shown on top. The y-axis states the −log 10 (p-value). The dotted horizontal line represents the threshold for Bonferroni correction (p=0.05/40).

We find that multiple upregulated clusters show enrichment for SCZ with the strongest evidence for the *synaptic function* cluster (MAGMA P=1.8×10^−7^, sLDSR P=7.2×10^−5^) (see Figure S7). For SRD, we find significant associations in the *transcription regulation* (P=2.5×10^−5^) and the *neuron formation* (P=1.2×10^−4^) gene cluster with MAGMA only. While the analysis of adult height using all differentially expressed genes did not yield any evidence for enrichment of genetic signal, enrichment is observed at the cluster level. The *cell connectivity* cluster (P=3.7×10^−4^) is enriched for height, in addition to suggestive enrichments in the *cell division* and *RNA regulation* cluster, which are not present for any of the psychiatric phenotypes. Remarkably, across all 8 clusters the enrichments of SCZ and height are inversely correlated (rho=−0.85, P=0.011, n=8, see also Figure S8).

Finally, in order to take into account the full spectrum of correlations and dependencies between clusters (Figure S9), we performed a conditional analysis for SCZ, the trait for which the strongest cluster enrichments are observed with both methods. Using the same MAGMA model, for each cluster, we conditioned on the highest gene members (membership > 0.5) of the other seven clusters (Table 2). We find that the SCZ enrichment is driven by the *synaptic function* cluster (p=2.88×10^−3^) only. The same conditional analysis for SRD, which only showed a significant enrichment with MAGMA, shows that this effect is primarily driven by the *transcription regulation* cluster (p=5.42×10^−3^) (Table S7).

**Table 2.** The association with SCZ risk is driven by the synaptic function gene cluster. Gene level association signal is regressed on cluster membership while adjusting for high membership genes of all other seven clusters. Shown are the results of the primary analysis (not adjusted for other clusters) and the conditional analysis with MAGMA. Beta = regression coefficient, SE = standard error.

| Schizophrenia - clusters | MAGMA Primary |  | MAGMA Conditional |  |
| --- | --- | --- | --- | --- |
|  | Beta (SE) | P-value | Beta (SE) | P-value |
| Cell division | -0.045 (0.017) | 1.00 | -0.047 (0.027) | 0.96 |
| RNA regulation | -0.040 (0.017) | 0.99 | -0.044 (0.027) | 0.95 |
| RNA processing | -0.006 (0.017) | 0.64 | -0.011 (0.024) | 0.68 |
| Neuron formation | 0.048 (0.017) | $2.12 \times 10^{-3}$ | 0.018 (0.036) | 0.30 |
| **Synaptic function** | **0.077 (0.017)** | **$1.82 \times 10^{-6}$** | **0.070 (0.026)** | **$2.88 \times 10^{-3}$** |
| Cell signaling | 0.052 (0.016) | $6.88 \times 10^{-4}$ | 0.032 (0.023) | 0.08 |
| Transcription regulation | 0.048 (0.016) | $1.67 \times 10^{-3}$ | 0.019 (0.025) | 0.22 |
| Cell connectivity | -0.061 (0.017) | 1.00 | -0.076 (0.026) | 1.00 |

## Discussion

We investigated a longitudinal *in vitro* stem cell model of human neuronal differentiation to study psychiatric disease susceptibility based on evidence from GWAS. Among five major psychiatric disorders, we observe that SCZ disease susceptibility is significantly enriched in a set of genes relevant to *synaptic functioning* that are upregulated during differentiation. We therefore propose *in vitro* human neuronal differentiation as an experimental system to further understand and decipher the polygenic architecture of psychiatric disease biology, in particular that of SCZ.

We confirmed that our *in vitro* model recapitulates neuronal signatures of *in vivo* cortical development across specific developmental time periods and laminae of the human neocortex. This is in line with previous findings^14^ and highlights that longitudinal gene expression dynamics underlying our model of human neuronal differentiation can be informative to study genes and pathways involved in *in vivo* human cortical development. Importantly, neuronal cell types^39, 40^ and early brain development^7, 22, 41^ have been postulated as integral components of SCZ disease susceptibility. Here, we observe that genes differentially expressed across neuronal differentiation are significantly associated with genome-wide disease risk of SCZ and that this risk aggregates in genes involved in synaptic functioning during development. Although not the only pathogenic process contributing to SCZ, synaptic dysfunction is most strongly supported by genetic data, postmortem expression studies, and animal models^40, 42–45^. We are the first to provide evidence for this hypothesis using a longitudinal *in vitro* cell-based model and aggregate polygenic disease risk. Two of the highest gene members of the *synaptic function* gene cluster enriched for SCZ include Calcium Voltage-Gated Channel Subunit Alpha 1C *(CACNA1C)* and Solute Carrier Family 45 Member 1 *(SLC45A1)*, both located at a genome-wide significant SCZ locus^22^. We find no evidence for AD, a late-onset non-psychiatric brain disease, nor for adult human height in this neuronal cluster. Together, our findings demonstrate that longitudinal transcriptomic signatures important for neuronal differentiation recapitulate the *in vivo* context *and* align with the genetic basis of the disease. SCZ disease biology, and in particular synaptic functioning, can thus be studied through these molecular processes captured by this *in vitro* model.

We also observed a significant enrichment of genetic signal with MAGMA for SRD in genes upregulated during differentiation, and show that this enrichment is predominantly driven by genes in the *transcription regulation* gene cluster. Interestingly, the SRD GWAS reported that the top SNPs were enriched for transcription regulation related to neurodevelopment^11^, which is in line with our *in vitro* findings. We observed no enrichment of the GWAS of recurrent and severe MDD in Han-Chinese women^23^. The latter sample represents the most genetically and phenotypically homogeneous GWAS of MDD. The fact that for these results no enrichment for any of our gene sets was observed may suggest that neurodevelopmental processes play a lesser role in MDD^46^. Alternatively, larger sample sizes are needed to better capture the genome-wide genetic risk associated with MDD. Self-reported depression is a much broader phenotype that may include other psychiatric traits, which could drive the observed neurodevelopment and transcription findings. Although it remains unclear how these results and the application of the model extrapolate to the MDD phenotype, our approach does highlight enrichment in distinct clusters for SRD and SCZ and could help shed light on how these two complex traits differ in their etiology.

We did not find any evidence of significant association in the neuronal clusters for ADHD. This could be due to the smaller sample size in the GWAS studies and thereby lack of power to find a significant association with our transcriptomic signatures (Figure S10). As GWAS sample sizes are expected to increase, these gene cluster associations should be revisited.

For height, we found effects in opposite direction of psychiatric traits in the downregulated gene clusters. Strikingly, we observe an inverse correlation between SCZ and height enrichment stratified across gene clusters (Supplementary Results, Figure S8 and S11), despite the absence of any evidence of a genetic correlation across the whole-genome^47^. These observations not only illustrate the added value of individual longitudinal gene clusters, but also highlight a complex genetic relationship between these two phenotypes.

A strength of our approach is the longitudinal analysis framework that we developed. We implemented an experimental design across a dense and repeatedly sampled time-series and integrated longitudinal transcriptomic signatures with genome-wide disease risk using available GWAS summary statistics. This increases statistical power to directly investigate the cumulative impact of risk loci on genes important to our model system. While we specifically chose to perform our experiments across an isogenic background to minimize variation and maximize statistical power to identify transcriptomic signatures, our framework can easily be extended to a multi-sample design (e.g. cases vs controls)^19, 48^, which makes it relevant for many disease-specific experimental settings.

Our experimental procedure applied differentiation towards a broad neuronal phenotype. Our work does not exclude disease associations with specific subtypes of neuronal cells nor with other major brain cell types. We provide a proof-of-concept of an *in vitro* model of neuronal cells for studying complex diseases, such as SCZ, and present an analytical framework that includes longitudinal assessment of gene expression profiles. This approach can readily be extended to study *in vitro* differentiation of other major brain cell types, such as astrocytes or oligodendrocytes. Although we show strong evidence for SCZ risk in early prenatal neurodevelopment, our findings do not preclude an additional contribution of postnatal neurodevelopment to the etiology of the disease^49–51^.

As GWAS risk loci have small effect sizes and are abundantly distributed across the genome, new approaches are needed that allow for functional investigation of polygenic disease architectures. Embracing the polygenic nature of psychiatric disorders is an important step forward in translating findings from GWAS to disease biology^52^. Our approach uses the cumulative impact of risk loci to identify gene clusters most strongly associated with SCZ while accounting for all other pathways identified in our model. This strategy allowed us to narrow down on potential core disease processes and opens up new avenues to study disease in the context of polygenicity. Future work can for example incorporate model perturbations to study aggregate disease risk in finer detail or use the model for functional fine-mapping of specific SCZ GWAS loci across an isogenic background in a controlled environment. Alternatively, cluster-specific polygenic risk scores (cPRS) can be used to study patients with a high genetic load versus those with a low genetic load to study *in vivo* or *in vitro* molecular and cellular differences. This will not only help elucidate the underlying disease mechanisms but can also provide insights into phenotypic domains within the heterogeneous schizophrenia patient population.

In summary, the current study establishes WA09 neuronal differentiation as an *in vitro* genomic tool to study psychiatric disease susceptibility. We demonstrate the value of an analytical framework that integrates longitudinal *in vitro* transcriptomic signatures with GWAS summary statistics to study the cumulative impact of risk loci on biological pathways. Overall, this work contributes to understand the functional mechanisms that underlie psychiatric disease heritability and polygenicity in the post GWAS era.

## Data Availability

The Illumina HT-12 v4 gene expression data is available through the Gene Expression Omnibus (GEO) archive *(Accession number is available for review).* This dataset has the raw and normalized gene expression values on all samples. Supplementary table 8 furthermore has specific probe annotations, such as probabilities of differential expression and probe membership values for all identified clusters.

## Author Contribution

The project was led by R.O. Experiments were designed and conceived by A.O. and R.O. Experiments were optimized, conducted and samples processed by A.O., M.B., and R.M. Analysis of the data was performed by A.O and M.B and feedback provided by L.O. and R.O. The main findings were interpreted by A.O., M.B., R.M., L.O., and R.O. Primary drafting of the manuscript was performed by A.O. and main feedback provided by R.O and L.O. All authors contributed to the production and approval of the final manuscript.

## Acknowledgement

We thank all research participants and researchers involved in making each GWAS summary statistic available and this work possible, including the 23andMe Research Team. We thank C. de Leeuw for his helpful input and troubleshooting with MAGMA analyses and thank the LD score regression team for their input and helpful troubleshooting with stratified LDSR. This research was supported by NIH/NIMH R01 MH090553.

